# The mechanical inhibition of the isolated V_o_ from V-ATPase for proton conductance

**DOI:** 10.1101/2020.02.13.946640

**Authors:** Jun-ichi Kishikawa, Atsuko Nakanishi, Aya Furuta, Takayuki Kato, Keiichi Namba, Masatada Tamakoshi, Kaoru Mitsuoka, Ken Yokoyama

## Abstract

V-ATPase is an energy converting enzyme, coupling ATP hydrolysis/synthesis in the hydrophilic V_1_ moiety, with proton flow through the V_o_ membrane moiety, via rotation of the central rotor complex relative to the surrounding stator apparatus. Upon dissociation from the V_1_ domain, the V_o_ of eukaryotic V-ATPase can adopt a physiologically relevant auto-inhibited form in which proton conductance through the V_o_ is prevented, however the molecular mechanism of this inhibition is not fully understood. Using cryo-electron microscopy, we determined the structure of both the *holo* V/A-ATPase and the isolated V_o_ at near-atomic resolution, respectively. These structures clarify how the isolated V_o_ adopts the auto-inhibited form and how the *holo* complex prevents the formation of this inhibited V_o_ form.

**One Sentence Summary:** Cryo-EM structures of rotary V-ATPase reveal the ON-OFF switching mechanism of H^+^ translocation in the V_o_ membrane domain.

## Main Text

Rotary ATPase/ATP synthases, roughly classified into F type and V type ATPase, are marvelous, tiny rotary machines (*1-5*). These rotary motor proteins share a basic molecular architecture composed of a central rotor complex and the surrounding stator apparatus. These proteins function to couple ATP hydrolysis/synthesis in the hydrophilic F_1_/V_1_ moiety with proton translocation through the membrane embedded hydrophobic F_o_/V_o_ moiety by rotation of the central rotor complex relative to surrounding stator apparatus, via a rotary catalytic mechanism (Figure 1) (*2-6*).

**Figure 1.**
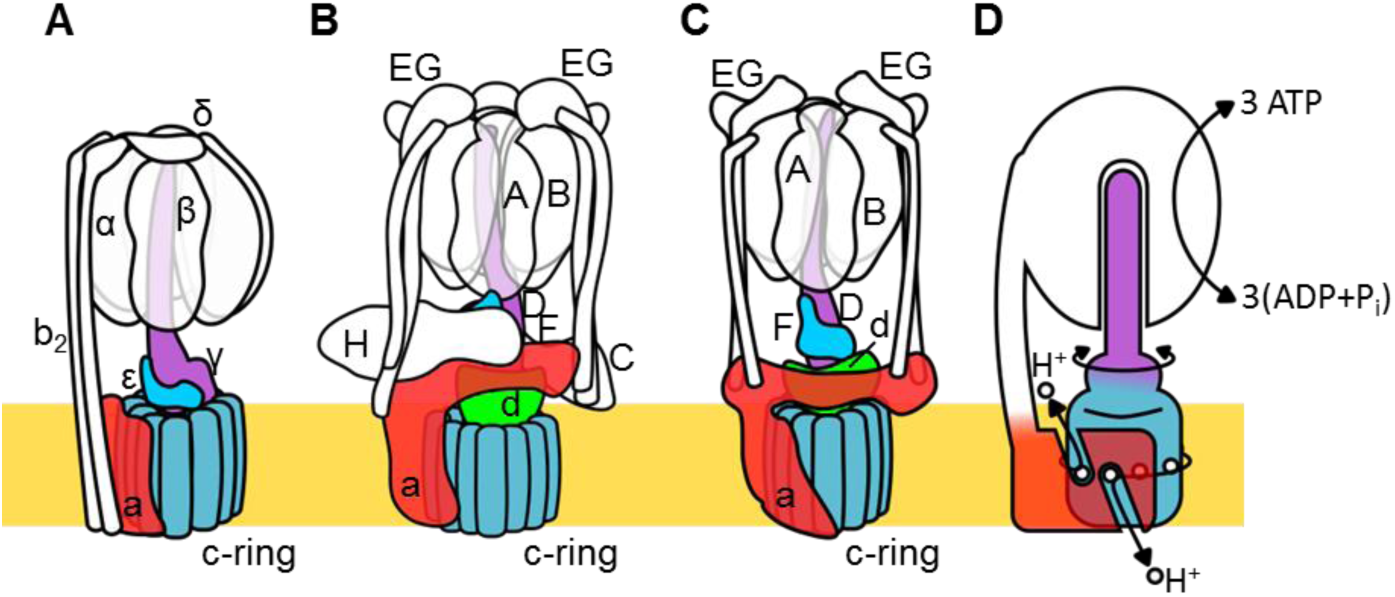
Schematic of rotary ATPase/synthases and the rotary catalytic mechanism. A. bacterial F_o_F_1_, B. yeast V-ATPase, C. *Tth* V/A-ATPase, D. schematic model of rotary catalytic mechanism. The subunits of the central rotor complex are colored: c-ring; dark blue, a-subunit; red, central axis; purple and cyan, and d-subunit; green.

Thus, both F and V type ATPases are capable of either ATP synthase coupled with proton motive force driven by membrane potential or proton pumping powered by ATP hydrolysis. F type ATPase (F-ATPase, or F_o_F_1_) in mitochondria functions as an ATP synthase coupled to respiration, whilst in some bacteria F-ATPase can function as an ATP dependent proton pump (*7, 8*).

V type ATPase (V-ATPase, or V_o_V_1_) resides mainly in the membranes of acidic vesicles in eukaryote cells, functioning as a proton pump using a rotary catalytic mechanism (*3, 9, 10*). Eukaryotic V-ATPases probably evolved from the prokaryotic enzymes (*11, 12*), which are termed Archaeal ATPase or V/A-ATPase (*3, 13*). V/A-ATPase from a thermophilic bacterium, *Thermus thermophilus* (*Tth* V/A-ATPase) is a rotary ATPase that has been well characterized using both structure and single molecular observation studies (*1, 9, 10, 14-18*). The overall structure of *Tth* V/A-ATPase closely resembles that of eukaryotic V-ATPase although it lacks some of the accessary subunits of the eukaryotic enzyme (Figure1B,C). The *Tth* V_1_ moiety is composed of four subunits with a stoichiometry of A_3_B_3_D_1_F_1_ and is responsible for ATP synthesis or hydrolysis (*19, 20*). Upon dissociation from V_o_, the isolated V_1_ shows only ATP hydrolysis activity accompanied by rotation of the DF shaft. The *Tth* V_o_ moiety, responsible for proton translocation across the membrane, contains a central rotor complex (*d*_1_*c*_12_) and stator apparatus made up of the *a* subunit and two EG peripheral stalks (*a*_1_E_2_G_2_). In *holo Tth* V/A-ATPase, proton motive force drives rotation of the *d*_1_*c*_12_ rotor complex relative to the surrounding stator, resulting in rotation of the entire central rotor complex (D_1_F_1_*d*_1_*c*_12_) and inducing sequential conformation changes in the A_3_B_3_ catalytic hexamer to produce three ATP molecules from ADP and inorganic phosphates per one rotation (Figure 1D).

Eukaryotic V-ATPase is regulated by a unique mechanism involving dissociation/association of V_1_, likely to be key in controlling the pH of acidic vesicles (*21-23*). In yeast, glucose depletion condition in the culture medium induces dissociation of V_1_ domain from V_o_ domain resulting in reduced proton pumping activity of V-ATPase (Figure S1A). It is likely that the dissociated V_o_ loses the ability to translocate protons as a result of auto-inhibition. In the structure of the dissociated V_o_ of yeast, the hydrophilic region of the *a* subunit (*a*_sol_) changes its conformation to prevent rotation of the rotor complex (*24, 25*). The yeast *a*_sol_ lies in close proximity to the *d* subunit, the rotor region of the isolated yeast V_o_ structure. Both the *a*_sol_ domain and the *d* subunit are hallmarks of the V-ATPase family and are lacking in F-ATPases (see Figure 1A-C) (*14*). Thus, the *a*_sol_ and the *d* subunit are probably to be key in auto-inhibition of the dissociated V_o_. However, the precise mechanism of V_o_ auto-inhibition, and thus prevention of proton leakage, is currently unknown.

Similar regulatory dissociation/association mechanism of V/A-ATPase in bacteria cells has not been reported, however, reconstitution experiments suggest an assembly pathway for the *holo* complex, in which the cytosolic V_1_ associates with the V_o_ in the membrane (Figure S1B)(*26*). Thus, proton leak through the V_o_ in the *Tth* membranes might be somehow also blocked by a similar autoinhibition mechanism to the eukaryotic enzyme. Indeed, the *Tth* V/A-ATPase and eukaryotic V-ATPase share very similar structures with both V_o_ moieties made up of the *a* and *d* subunits in addition to the c ring.

Structural analysis using cryogenic microscopy (cryoEM) of the *holo* V/A-ATPase, including our recent study, revealed several rotational states of the entire *holo* complex (*17, 27*). However, understanding of the inhibition mechanism of the isolated *Tth* V_o_ is currently limited due to a lack of a high resolution structure.

Here, we report a cryoEM structure of isolated *Tth* V_o_ at 3.9 Å resolution. Our results clarify the molecular mechanism of proton leak inhibition from *Tth* cells through an assembly intermediate V_o_ of *holo* V/A-ATPase under physiological conditions.

### CryoEM structures of the isolated V_o_ and *holo Tth* V/A-ATPase

We purified both *Tth* V/A-ATPase and V_o_ via a His_3_-tagged *c* subunit from membranes of *T. thermophilus* cells using Ni-NTA resin. For *Tth* V/A-ATPase, acquisition of micrographs was carried out using a Titan Krios equipped with a Falcon II direct electron detector. Cryo-EM micrographs of the complexes reconstituted into nanodiscs resulted in higher resolution EM maps compared with the LMNG solubilized preparation previous reported (*17*). The strategy of single particle analysis for *Tth* V/A-ATPase is summarized in Figure S2A. The final structure of state 1 has an overall resolution of 3.6 Å (Figure 2A). After subtraction of the EM density of the membrane embedded domain from the density of the whole complex, we obtained a focused density map of A_3_B_3_D_1_F_1_*d*_1_ with two EG peripheral stalks and the soluble arm domain of the *a* subunit (*a*_sol_) at 3.5 Å resolution. This map allowed us to build an atomic model of A_3_B_3_D_1_F_1_ (V_1_). In our map, the obvious density of ADP-Mg was observed in the closed catalytic site, but not clearly observed in semi-closed site, in contrast to our previous structure of state 1 (5Y5Y). The secondary ADP in the semi-closed site shows lower occupancy, it is due to the low affinity of the semi-closed site for nucleotide and partial flexibility in the complex (Figure S3A). In the recent cryoEM map of *Tth* V/A-ATPase (6QUM), clear densities likely to correspond to ADP were observed in the cavities of the crown-like structure formed by the six β barrel domains of A_3_B_3_ (*27*). In contrast, these densities were not clearly visible in our structure (Figure S3B). These differences are presumably due to the purification procedures; we purified the His-tagged *Tth* V/A-ATPase using a nickel column, while the authors of the other study isolated their *Tth* V/A-ATPase without affinity purification.

**Figure 2.**
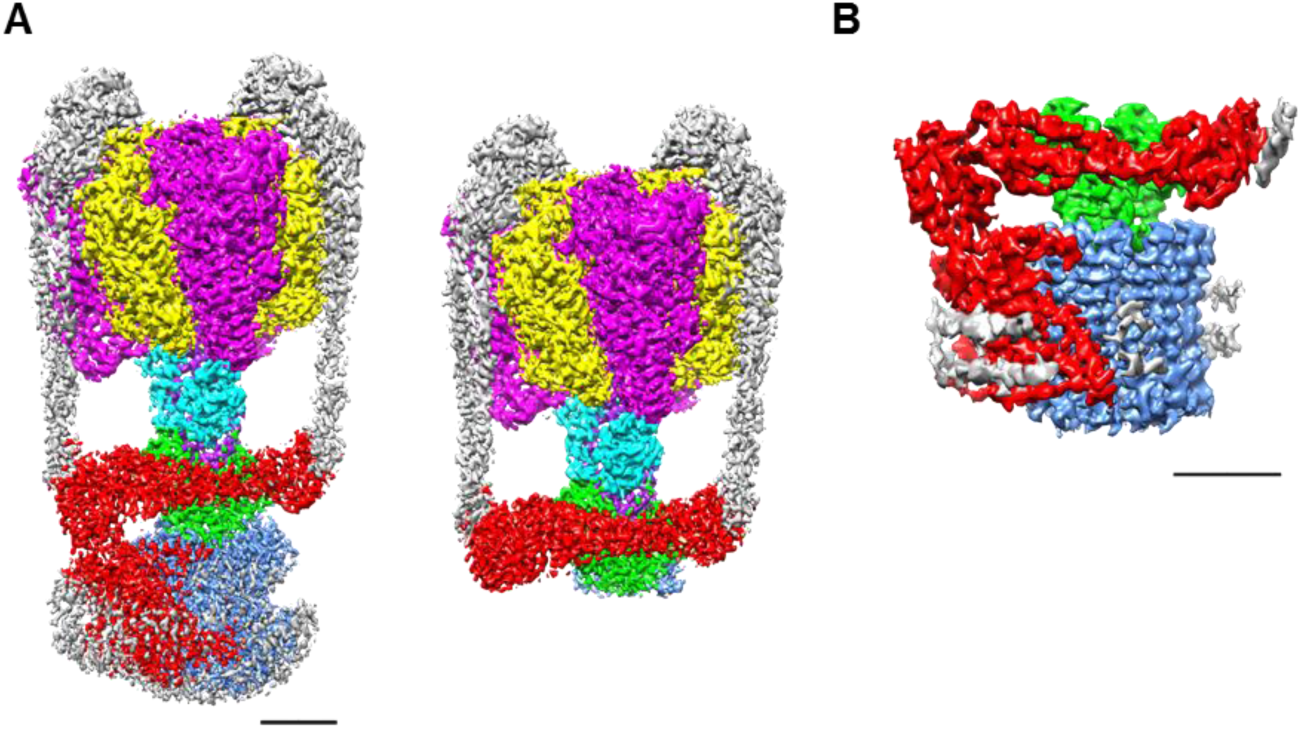
EM density map of complex. A. *holo Tth* V/A-ATPase (left) and focused refined map of A_3_B_3_DF*d*(EG)_2_*a*_sol_ (right) B. the isolated V_o_ (B). The density corresponding each subunit is colored: A; magenta, B; yellow, D; purple, F; cyan, E and G; gray, *a*; red, *d*; green, and *c*; dark blue. Scale bar; 30 Å.

Purified V_o_ reconstituted into nanodiscs was subjected to single particle analysis using a cryoEM (CRYOARM200, JEOL) equipped with a K2 summit electron direct detector in electron counting mode. The 2D class averages showed the isolated V_o_ with clearly visible transmembrane helices and a hydrophilic domain extending above the integral membrane region (Figure S2C). The density for the scaffold proteins and lipids of the nanodiscs is clearly visible surrounding the membrane domain of the isolated V_o_. Following 3D classification of the V_o_, only one major class was identified indicating that the isolated V_o_ is very structurally homogenous, in contrast to the *Tth* V/A-ATPase which is clearly visible in three different rotational states (*17*). Our 3D reconstruction map of the isolated V_o_ was obtained with an overall resolution of 3.9 Å. The final map shows clear density for protein components of V_o_, including subunit *a, d, c*_12_ ring, but the EM density for both EG stalks, which attach to the *a*_sol,_ is weak indicating disorder in these regions, suggesting their flexibility (Figure 2B). In this structure, the C-terminal region of the EG stalk on the distal side is visible. With the exception of these two EG stalks, side-chain densities were visible for most of the proteins in the complex, allowing construction of a *de novo* atomic model using phenix and coot (Figure 3A,B). The map contains an apparent density inside the *c*_12_ rotor ring, likely corresponding to the phospholipids capping the hole of the ring (Figure S4A). A further apparent density was identified in the cavity between the *a* subunit and the *c*_12_ ring on the upper periplasmic side (Figure S4B). This also might be corresponded to phospholipid and we postulate that this functions to plug the cavity between the *a* subunit and the *c*_12_ ring preventing proton leak from the periplasmic proton pathway. The densities corresponding to these phospholipids in our V_o_ structure are also observed in recently published cryoEM density map of the *holo* complex (*27*). Notably, the diameter of the *c*_12_ rotor ring in the isolated V_o_ is slightly smaller than that in the *Tth* V/A-ATPase (Figure S5A). It is likely that penetration of the short helix of the subunit D into the cavity of subunit *d* enlarges the diameter of the *c*_12_ rotor ring in the *Tth* V/A-ATPase.

**Figure 3.**
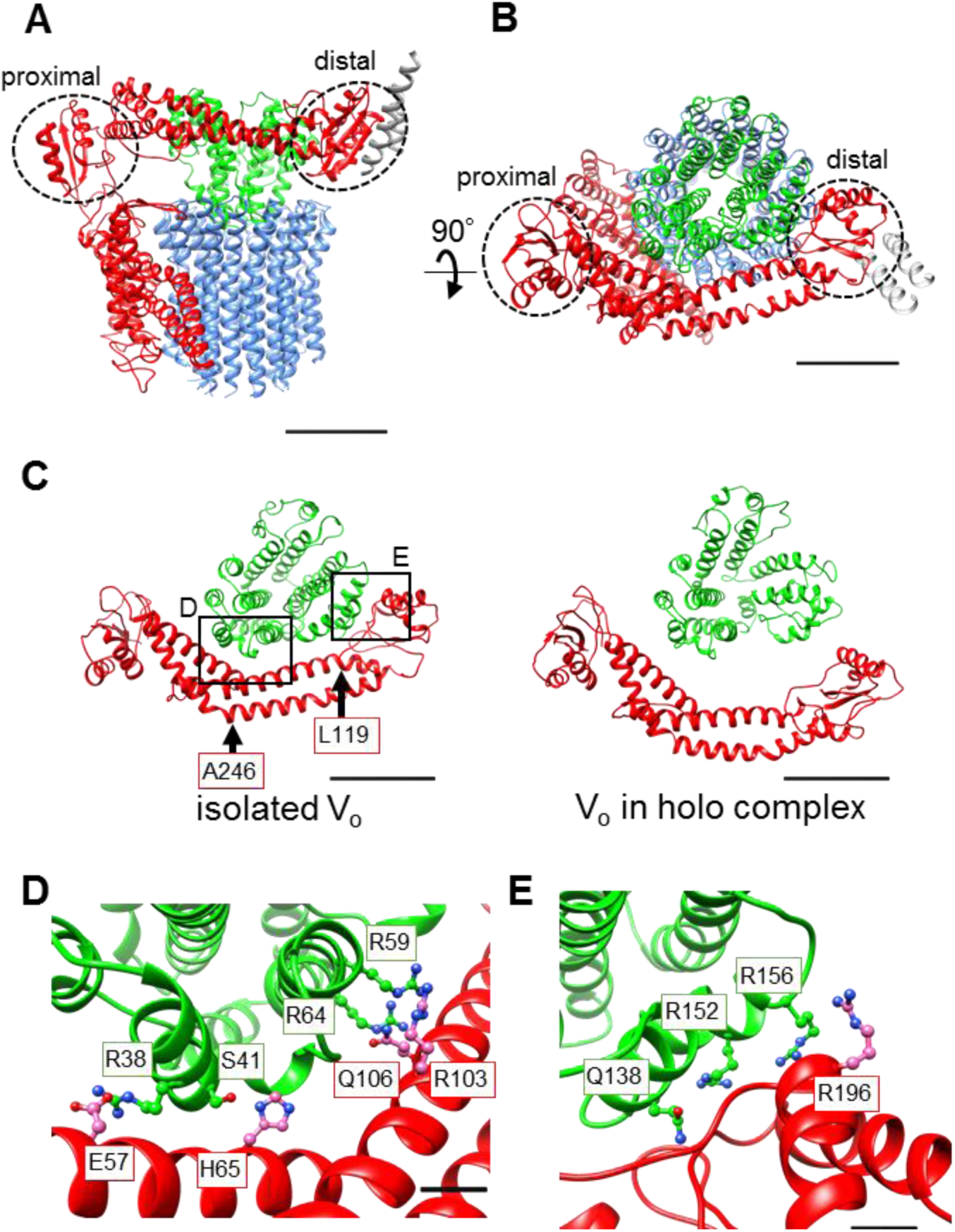
Atomic model of the isolated V_o_. A. Side view and B. Upper view of *a*-, *d*-, *c*-, and EG subunits colored as in Figure 2, respectively. Scale bar represents 30 Å. The proximal and distal subdomains of *a*-subunit are circled by the dotted lines. C. Comparison of the relative positions of *a*_sol_ (red) and the d subunit (green) in the isolated V_o_ (left) and the V_o_ moiety in the *holo* complex (right). Arrows indicate the kinking and twisting points in the *a*_sol_ in isolated V_o._ Scale bar represents 30 Å. D, E. Specific interactions between the *a*_sol_ and *d* subunit at proximal (D) and distal (E) regions. The regions are specified in black squares in C. The residues are represented as balls and sticks. Scale bar; 5 Å.

### Structure comparison of the isolated V_o_ with the *holo* complex

A comparison of our structure of the isolated V_o_ with that of V_o_ moiety in *holo* complex revealed a high degree of similarity in the membrane embedded region. However, there were significant differences in the *a* subunit. The basic structure of the *a* subunit of *Tth* V_o_ is almost identical to the eukaryotic counterpart, with both composed of a soluble arm domain (*a*_sol_) and a C-terminal hydrophobic domain responsible for proton translocation via rotation of the *c*_12_ ring. The *a*_sol_ contains two globular α/β folding subdomains responsible for binding of both the proximal and distal EG stalks (Figure 3A and B). Both globular subdomains are connected by a hydrophilic coiled coil with a bent conformation.

In contrast to the structure of V_o_ moiety in the *holo* complex, the *a*_sol_ in V_o_ only is in close proximity to the *d* subunit as a result of kinking and twisting of the coiled coil at residues *a*/L119 and *a*/A246 (Figure 3C, indicated by the arrows). In this structure, there are several interactions between the residues in the *a*_sol_ and the *d* subunit (Figure 3D). At the proximal site, three amino acid residues, *a*/E57, *a* /H65, and *a*/Q106, form salt bridges or hydrogen bonds with residues *d*/R38, *d*/S41, and *d*/R64 in the *d* subunit, respectively. The side chain of *d*/R59 likely forms π-π stacking with *a*/R103. Our structure also revealed clear connected densities between the distal subdomain of the *a*_sol_ and the *d* subunit (Figure 3E). Four side chains, *d*/Q138, *d*/R152, *d*/R156, and *a*/R196 probably form hydrogen bonds with the oxygen atoms in the main chain of *a*/E201, *a*/L144, *a*/A197, and d/R156, respectively. With the exception of the interaction between *a*/E57 and *d*/R38 in the proximal site, these interactions are broken by the dynamic movement of the *a*_sol_ and conformational change of *d* subunit in the V_o_ moiety of *holo Tth* V/A-ATPase. These conformational changes of the isolated V_o_ induced by binding of V_1_ (A_3_B_3_DF) to the V_o_ are described in a separate section below.

### Voltage threshold for proton conductance activity of the isolated V_o_

Our structure of the isolated V_o_ suggests that the rotation of *c*_12_ rotor ring relative to the stator is mechanically hindered by a defined interaction between the *a*_sol_ and *d* subunit. To investigate this mechanical hindrance of proton conductance through the V_o_, we reconstituted the isolated V_o_ into liposomes energized with a *Δψ* generated through a potassium ion (K^+^)/valinomycine diffusion potential. The pH change in the liposomes was monitored with 9-Amino-6-Chloro-2-Methoxyacridine (ACMA); the emission traces at 510 nm excited at 460 nm were recorded (Figure 4). The size of the membrane potential was modulated by varying the external K^+^ concentration. As shown in Figure 4B, a voltage threshold was observed in that the isolated V_o_ shows no proton conductance at less than 120 mV of membrane potential. When the membrane potential is 130 mV or more, the proton conductance through the V_o_ increases in proportion to the membrane potential (Figure 4B). The reported membrane potential in bacteria cells is - 75 ∼ -140 mV (*28*). Thus, the observed inhibitory mechanism of the isolated V_o_ can function to prevent proton leak through the V_o_ under physiological conditions. In contrast to the V_o_, several experiments have indicated that proton conductance through F_o_ of bacteria does not show the threshold of membrane potential (*29*). Together, the observed results strongly suggest that the *a*_sol_ of the *a* subunit and the *d* subunit, absent in F_o_ and hallmarks structure of the V type ATPases, are key for mechanical inhibition of proton conductance through V_o_.

**Figure 4.**
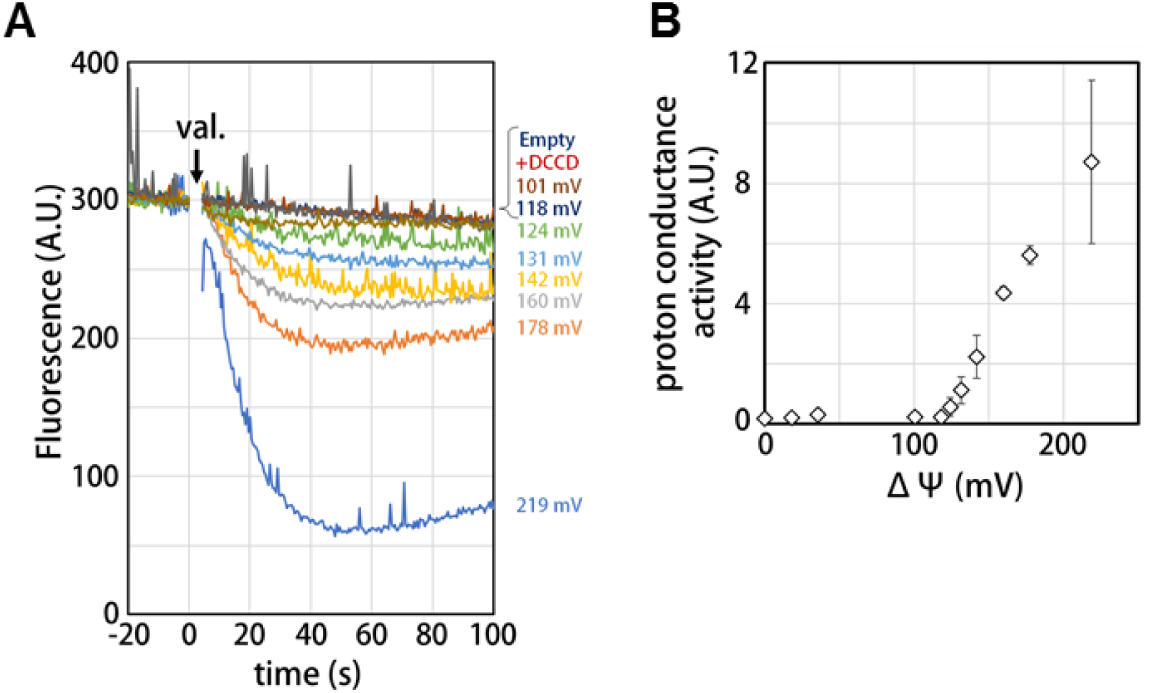
Proton conductance through the isolated V_o_. A. Changes of fluorescence of ACMA due to pH changes inside the V_o_ proteo-liposomes. Membrane potential (ΔΨ) values were estimated by the Nernst equation; ΔΨ=*RF*/*zF* ln[KCl]_o_/[KCl]_i_, described in the Methods section. B. Voltage threshold of the proton conductance through the V_o_.

### Structure of the membrane embedded region of the isolated V_o_

Our atomic model of V_o_ presented here reveals details of both proton paths formed by the membrane embedded C-terminal region of the *a* subunit (*a*_CT_) and its interface with the *c*_12_ ring. The *a*_CT_ contains eight membrane embedded helices, MH1 to MH8. MH7 and MH8 are highly tilted membrane embedded helices characteristic of rotary ATPases. The cytoplasmic hydrophilic cavity is formed by the cytoplasmic side of MH4, MH5, MH7, and MH8, and the *c* subunit /chainZ. The cavity is lined by polar residues, *a*/R482, *a*/H491, *a*/H494, *a*/E497, *a*/Y501, *a*/E550, *a*/Q554, *a*/T553, *a*/H557, and *c*(Z)/Thr54 (Figure 5A), which seem to make up the cytoplasmic proton path. The periplasmic sides of MH1, MH2, MH7 and MH8 form the periplasmic hydrophilic cavity, lined with *a*/D365, *a*/Y368, *a*/E426, *a*/H452, *a*/R453, *a*/D455, and *c*(Y)/E63. The two hydrophilic channels are separated by a salt bridge formed between *c*(Z)/63Glu, a residue critical for proton translocation, and *a*/Arg563, *a*/Arg622 and *a*/Gln619 of MH7 (Figure 5B). This salt bridge is conserved in both eukaryotic and prokaryotic V_o_ (25,26). In contrast the salt bridge forms between a single arginine residue and a single glutamic (or aspartic) acid residue in F_o_(*5, 30, 31*). Similar to the two channel model described for other rotary ATPases (*32, 33*), the two arginine residues on the MH7 and 8 play an important role in protonation and deprotonation of the carboxy groups on the *c*_12_ ring, with the resulting rotation of *dc*_12_ driven by proton translocation from periplasmic to cytoplasmic sides. Notably, in addition to the rigid salt bridge formed between the two *a*/Arg residues, *a*/Gln and *c*/Glu, interactions between the *a*_ct_ and *c*_12_ ring are observed; *a*/Asp392 and Leu393 - *c*(Y)/Arg49 in the loop region of the *c* subunit (Figure S6A), and the periplasmic sides of MH5 and MH6 are in close proximity to the C-terminal end of the *c* subunit (Figure S6B). Overall, our V_o_ structure is largely identical to the V_o_ moiety in *holo* complex with the exception of key alterations in hydrophilic domain (*27*).

**Figure 5.**
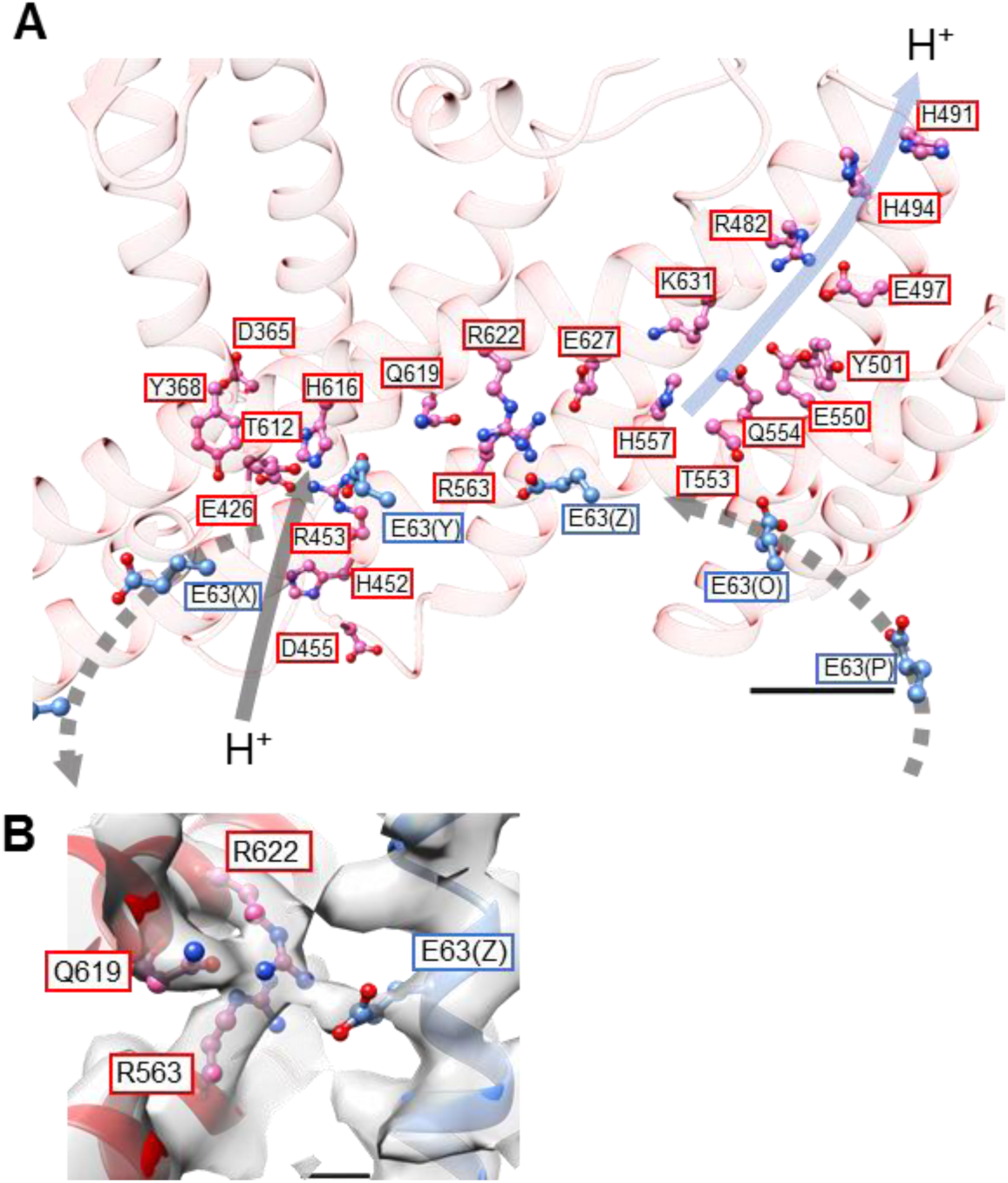
Structure of the hydrophobic domain of the isolated V_o_. A. Proton paths on both the cytoplasmic and periplasmic sides of the isolated V_o_. Residues lining the paths are represented as balls and sticks. Residues from the *a*-subunit and *c*-subunit are indicated in the red and blue boxes, respectively. Proton flow from the periplasmic side is represented by the grey arrow as it would occur in the case of ATP synthesis. Scale bar; 10 Å. B. Salt bridge between *a*/Arg563, Arg622, Gln619 and *c*/Glu63. Scale bar; 3 Å.

### Molecular basis of the auto-inhibition of proton conductance in the isolated V_o_

The inhibition mechanism of V_o_ depends upon conformational changes in two subunits. In the isolated V_o_, the *d* subunit adopts the closed form in which three side chains of the *d* subunit are able to interact with the distal subdomain of *a*_sol_. Once the short helix of the D subunit inserts into the cavity of the *d* subunit, the interaction between H6 and H11 via *d*/R90 and *d*/E195 is broken (Figure 6A and Movie S1), resulting in the *d* subunit adopting an open form where the orientation of three side chains move away from the distal subdomain of *a*_sol_.

**Figure 6.**
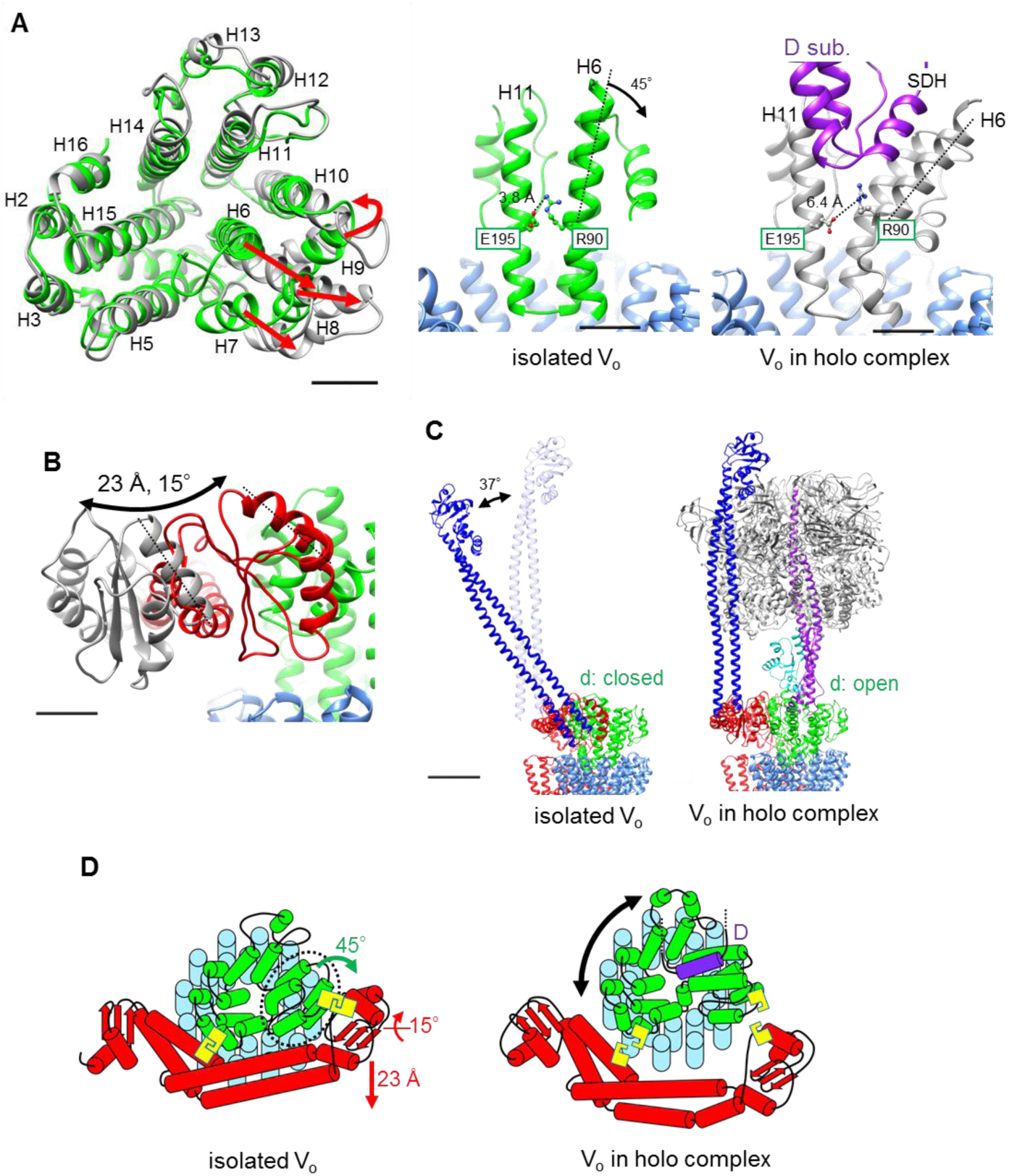
Conformational changes occurring in both the *d*- and *a*_sol_ subunits as a result of binding of V_1_ to V_o_. A. Structural changes in the *d* subunit caused by insertion of the screw driver helix (SDH). (left) Top view of *d*-subunit. The *d*-subunit from the isolated V_o_ and the *holo* enzyme are colored in green and grey, respectively. Red arrows indicate the movements of helices 6-9 (H6-9). (center, right) Key helices (H6 and 11) of *d*-subunit in the isolated V_o_ and the *holo* complex. The H6 bends 45° as a result of binding of SDH of D-subunit. B. Structural change of the distal subdomain of *a*_sol_. Upon the pivoting movement of *a*_sol_ on the proximal subdomain, the distal subdomain swings 25 Å and twist 15° between the isolated V_o_ (red) and the *holo* complex (gray). C. EG structure in the distal subdomain of *a*_sol_ (EG_d_) in the isolated V_o_ (left) and in the *holo* complex (right). D. Schematic representation of the mechanical inhibition of the V_o_ induced by dissociation of V_1_. In isolated V_o_, the rotation of central rotor is inhibited by interactions between *d*- and *a*_sol_ (yellow box, Figure 3D, E).

Another contributing factor is dynamic motion of the *a*_sol_ induced by binding of the distal EG stalk to the top of the A_3_B_3_. In the isolated V_o_, the C-terminal region of the EG stalk binding onto the distal subdomain of *a*_sol_ is at a much steeper angle relative to the horizontal coiled coil structure of *a*_sol_ than that in the *holo* enzyme (Figure 6B, C and S7). Once the N-terminal globular domain of the distal EG stalk binds onto the top of A_3_B_3_, the angled distal EG adopts a vertical standing form, resulting in both a twisting and kinking of the coiled coil of the hydrophilic arm and the distal globular subdomain (Figure 6C, Movie S2). These dynamic motions of the *a*_sol_ of *a* subunit induces disruption of the specific interactions of *a*_sol_ with *d* subunit.

The isolated yeast V_o_ also adopts a similar inhibited conformation where the *a*_sol_ is in close proximity to the *d* subunit, resulting in interaction between the stator and the rotor and inhibition of proton conductance (*24, 25*). Although an atomic model of yeast *holo* V-ATPase has yet to be determined, the *a*_sol_ is some distance from the *d* subunit in the V_o_ moiety of the poly alanine model of yeast V-ATPase (*34*). These structures hint at a similar conformational change in V_o_ induced by binding of the V_1_ domain as predicted in the *Tth* V/A-ATPase. Notably, the *d* subunit in the yeast *holo* complex adopts the open form, in contrast to the *Tth* V_o_ where the *d* subunit is in the closed form (Figure S8). With this single exception, the eukaryotic and prokaryotic V-ATPases seem to share a similar auto-inhibited mechanism of V_o_ preventing proton leakage from cells or acidic vesicles. This suggests that the auto-inhibition mechanism of V_o_ is conserved during the evolution of V type ATPases.

The interaction between the *a*_sol_ and *d* subunit stabilizes the isolated V_o_ structure and protects against loss of *d*-subunit in the absence of the rotor-stator interactions mediated by V_1_ as a result of the dissociation of the two domains (*35*). This stabilization of V_o_ is most likely to be key for both assembly of *holo* V-type ATPase complexes and regulation of eukaryotic V-ATPase via dissociation of V_1_ from V_o_.

## Acknowledgements

We are grateful to all the members of the Yokoyama Lab for their continuous support and technical assistance. Our research was supported by Grant-in-Aid for Scientific Research (JSPS KAKENHI) Grant Number 17H03648 to K.Y. Our research was also supported by Platform Project for Supporting Drug Discovery and Life Science Research (Basis for Supporting Innovative Drug Discovery and Life Science Research (BINDS)) from AMED under Grant Number JP17am0101001 (support number 1312), and Grants-in-Aid from “Nanotechnology Platform” of the Ministry of Education, Culture, Sports, Science and Technology (MEXT) to K.M. (Project Number. 12024046)

## Author contributions

JK, and AN designed, performed and analyzed the experiments. JK, AN, AF, TK, and KM analyzed the data and contributed to the preparation of the figures. MT constructed vectors for expression of mutant proteins. TK, and KM provided technical support and conceptual advice. KY designed and supervised the experiments and wrote the manuscript. All authors discussed the results and commented on the manuscript.

## Data and materials availability

The density maps and the built models for *Tth* V_o_V_1_, *Tth* V_1_ (focused refined), and *Tth* V_o_ were deposited in EMDB (EMDB DI; 30013, 30014, and 30015) and PDB (PDB ID; 6LY8 for V_1_ and 6LY9 for isolated V_o_), respectively. All data is available in the main text or the supplementary materials.

## Supplementaly Materials

Materials Methods

Figure S1-S10

Tables S1

Movies S1 and S2

References (36-47)

